# Predictors of arm non-use in chronic stroke: a preliminary investigation

**DOI:** 10.1101/702159

**Authors:** Laurel J. Buxbaum, Rini Varghese, Harrison Stoll, Carolee J. Winstein

## Abstract

**Background:** The phenomenon of non-use after stroke is characterized by failure to use the contralesional arm despite adequate capacity. It has been suggested that non-use is a consequence of the greater effort and/or attention required to use the affected limb, but such accounts have not been tested, and we have poor understanding of the characteristics of individuals who exhibit non-use.

**Objective:** We aimed to provide preliminary evidence regarding the demographic, neuropsychological, and psychological factors that may influence non-use in chronic stroke.

**Methods:** Twenty chronic stroke survivors (10 left and 10 right hemisphere stroke) with mild-to-moderate sensory-motor impairment on the upper extremity Fugl-Meyer (UEFM) were assessed with the Actual Amount of Use Test (AAUT), which measures the disparity between amount of use and quality of movement in “spontaneous” versus “forced” conditions. Participants were als assessed with measures of limb apraxia, spatial neglect, attention/arousal, and self-efficacy (confidence in arm movement). Using stepwise multiple regression, we determined whether demographic information and/or performance on these measures predicted AAUT non-use scores.

**Results:** Scores on the UEFM as well as attention/arousal and self-efficacy predicted the degree of non-use. Attention/arousal predicted non-use above and beyond UEFM.

**Conclusions:** Given the complexity of the non-use phenomenon, it follows that a combination of impairment, attention/arousal, and perceived confidence predicted non-use behavior. That a measure of attention/arousal predicted non-use behavior above and beyond sensory-motor functioning highlights the importance of motivated engagement to drive use of the paretic limb. Larger-scale studies incorporating additional measures (e.g., mental health, lesion volume and white matter connectivity, pain, motivation) will be important for future investigations.

## Introduction

One of most vexing problems in neurorehabilitation is the phenomenon of arm non-use. After a stroke, the contralesional arm may exhibit functional motor and sensory impairments that include muscle weakness and reduced proprioception and tactile sensation. These deficits lead to immobility, which is in turn related to spasticity, stiffness, pain, and abnormal reflex mechanisms.^1^ As a further consequence, many patients exhibit reduced use of the contralesional arm, whether via a maladaptive learning process in which movement is suppressed over time ^2^ or via a more direct stroke-related reduction in volitional drive toward limb use.^3^

Irrespective of its precise underlying mechanism from a physiological perspective, arm non-use is likely to negatively influence clinical and research rehabilitation efforts. It is frequently assumed that measured improvements in capacity as a result of training translate to increased use of the arm in daily activities.^4^ However, recent evidence suggests that despite gains in capacity as a result of rehabilitation, there may be little to no improvement in use of the arm in daily life. For example, a secondary analysis from a Phase II dose-response trial conducted at Washington University aimed to assess changes in the performance of 78 participants with chronic stroke during daily activities over an 8-week intervention period. The investigators used accelerometers to assess overall arm use, movement intensity, or acceleration parameters at weekly intervals. Neither changes in capacity, as measured by the Action Research Arm Test, nor overall amount of practice influenced the amount of use as measured by accelerometer variables.^4^ Baseline arm capacity, stroke chronicity, concordance (dominant side = affected side), and ADL status all influenced the intercepts but not the slopes of these variables. These data reinforce the importance of assessing capacity and performance separately (see ^5,6^). Further, Stewart & Cramer (2013) highlight the importance of using patient-reported measures for providing insights into motor function after stroke.^7^ This observation influenced the choice of measures used here.

Non-use was the impetus for the development of constraint-induced movement therapy (CIMT), an intervention that combines restraint of the unaffected upper extremity with forced training of the affected extremity using a shaping protocol. CIMT in chronic stroke shows evidence of efficacy in improving arm function as compared to traditional rehabilitation.^8–13^ A recent meta-analysis of studies conducted with acute and post-acute stroke participants^14^ showed evidence supporting improved arm function from CIMT. However, there was inconsistent evidence for improvements in actual amount of use in daily activities (as measured by the Motor Activity Log).

Initial development of CIMT was based on a model of conditioning, informed by the observation that monkeys prevented from experiencing early failures did not develop non-use.^15^ On this model, CIMT achieved via an extrinsic factor—a constraint or mitt—extinguishes learned non-use. Alternative models, however, suggest that non-use is a consequence of the greater effort (e.g., force or distance commands) and/or attention (e.g., to proprioceptive, somatosensory, and visual feedback) required to use the affected limb. Spontaneous use may therefore reflect a trade-off between effort and disability, wherein the unaffected limb will be used alone whenever this may lead to a reasonably successful outcome.^16^ On this account, and based on models that emphasize intrinsic motivational factors (e.g., OPTIMAL^17^), overcoming non-use requires that we address the motivational components of the disorder. In support of this possibility, CIMT with a “transfer package” including a behavioral agreement, diary use, problem-solving to overcome perceived barriers, home skill practice assignments, and weekly telephone calls has shown efficacy superior to CIMT alone for performance-based behavioral ^12,18^ and structural brain changes.^19^

An important step in the development of targeted treatments for failure to use the limb despite adequate functional ability is a better understanding of the characteristics of individuals who exhibit this phenomenon. In this preliminary study we carefully selected a representative sample of individuals with chronic stroke who exhibited a range of sensory-motor impairments to begin to develop hypotheses about the demographic, neuropsychological, and psychological factors that may influence non-use behavior.

## Methods

### Participants

Participants were 20 individuals with chronic (> 12 months post) stroke, 10 of whom had suffered a single left hemisphere stroke (LCVA) and 10 of whom had experienced a single right hemisphere stroke (RCVA). Table 1 provides demographic information. Mean age was 62.5 years, range 48-75 years; mean education was 14.3 years, range 12-23 yrs; mean chronicity was 5.8 years, range 1.1-13.1 yrs; and mean Fugl-Meyer was 46, range 30-63. Five LCVA and 5 RCVA were recruited from the Neurocognitive Rehabilitation Research Registry at Moss Rehabilitation Research Institute;^20^ the remaining 5 LCVA and 5 RCVA were recruited from the Registry for Aging and Rehabilitation Evaluation database of the Motor Behavior and Neurorehabilitation Laboratory at the University of Southern California. All were right-handed and achieved a score on the Upper Extremity Fugl-Meyer (UEFM) of at least 29, demonstrating moderate to mild motor impairment.^21,22^ Patients with a history of psychosis, neurologic disorder, traumatic brain injury, or alcohol/drug abuse were excluded. Participants at the two sites were matched for age (t(18) = .06, p = .95), chronicity (t(18) = -.41, p = .68, and UEFM scores (t(18) = .56, p = .58); the USC sample tended to be slightly more highly educated than the Moss sample (t(18) = −1.99, p = .06). Similarly, LCVA and RCVA were well matched in age, chronicity, education, and UEFM scores (t(18) < 1.2, p >. 23, for all comparisons). The Moss and USC protocols were approved by the Institutional Review Boards of Einstein Healthcare Network and University of Southern California, respectively. Participants at Moss were paid for their participation in the study consistent with Moss policy.

**Table 1.** Demographic information.

| **Subject** | **Site** | **Hemisphere**<br>**of stroke** | **Age**<br>**(yrs)** | **Education**<br>**(yrs)** | **Years**<br>**post-**<br>**stroke** | **Fugl-**<br>**Meyer**<br>**Score** |
| --- | --- | --- | --- | --- | --- | --- |
| S1 | USC | Left | 56 | 16 | 7.08 | 30 |
| S2 | USC | Left | 48 | 12 | 4.76 | 37 |
| S3 | USC | Left | 58 | 14 | 3.08 | 52 |
| S4 | USC | Right | 63 | 14 | 3.27 | 36 |
| S5 | USC | Right | 73 | 16 | 12.04 | 39 |
| S6 | USC | Left | 50 | 18 | 5.52 | 51 |
| S7 | USC | Right | 72 | 16 | 6.62 | 47 |
| S8 | USC | Left | 71 | 12 | 2.19 | 55 |
| S9 | USC | Right | 66 | 13 | 9.29 | 51 |
| S10 | USC | Right | 62 | 23 | 7.54 | 54 |
| S11 | MRRI | Left | 60 | 14 | 6.17 | 39 |
| S12 | MRRI | Left | 71 | 13 | 13.58 | 49 |
| S13 | MRRI | Left | 52 | 12 | 4.33 | 50 |
| S14 | MRRI | Left | 75 | 13 | 4.92 | 51 |
| S15 | MRRI | Left | 67 | 12 | 1.08 | 63 |
| S16 | MRRI | Right | 52 | 13 | 5.33 | 34 |
| S17 | MRRI | Right | 68 | 14 | 8.17 | 47 |
| S18 | MRRI | Right | 56 | 13 | 1.5 | 49 |
| S19 | MRRI | Right | 64 | 12 | 7.17 | 41 |
| S20 | MRRI | Right | 62 | 16 | 2.92 | 50 |

### Background Test Battery

Background tests included a sample of neuropsychological (e.g., apraxia), and psychological predictors including perceived confidence.

#### Apraxia task (Imitation of novel gestures)

Prior investigations indicate that limb apraxia (spatio-temporal deficits in imitation, pantomime of tool use movements, and/or tool use, even with the ipsilesional limb) is a potent predictor of disability after left hemisphere stroke.^e.g.,23^ To assess spatio-temporal deficits without the confound of a tool knowledge component, we used our laboratory’s well-studied assessment of meaningless gesture imitation. Participants were shown videos of an experimenter performing 10 novel gestures while facing the viewer. The gestures were meaningless analogues of meaningful tool use gestures (see ^24^ for details of stimulus development). There were two versions of the videos, and both LCVA and RCVA patients used their ipsilesional hand to copy the model in mirror perspective. Each gesture was shown twice in succession, and participants were permitted to begin imitating while observing. Participants’ gestures were videotaped and later scored using scoring criteria long in use at Moss (see ^25^). Both USC and Moss patients’ performance was scored by trained coders in the Buxbaum lab who demonstrated reliability with previous coders in the lab, as defined by Cohen’s Kappa >0.85 (‘very good’ agreement ^26^).

#### Spatial neglect and attention/arousal task

The lateralized and non-lateralized attention/arousal deficits that are the core components of the spatial neglect syndrome are strong predictors of outcome in stroke.^e.g.,27^ To assess both lateralized and non-lateralized attention, participants performed the Virtual Reality Lateralized Attention Test (VRLAT), short form. In this game-like task, participants use a computer joystick to navigate down a winding path on a computer monitor. Images of a variety of static objects including colored trees, animal statues, and road signs are observable in quasi-randomized locations on each side of the path; moving objects including balls and skateboards cross the path at random intervals, and there are random distracting noises. Participants’ task is to name the color of the trees and the animals depicted by the statues. The course is traversed once “coming” and once “going”, so the same objects appear once on each side of the path. Points are given for objects correctly named with a full description (“purple tree”, “cat statue”). The normalized difference in points awarded to the ipsilesional and contralesional sides (ipsi – contra /(ipsi + contra) is a measure of spatial neglect. Following from prior research,^e.g.,28^ total points irrespective of side (ipsi + contra) served as a measure of non-lateralized attention and arousal. ^29^

#### Self-efficacy questionnaire

We used the 20-item Confidence in Arm and Hand Movement (CAHM) questionnaire to evaluate perceived self-efficacy in performing unimanual and bimanual functional tasks using the contralesional upper extremity in home and community contexts in patients with stroke (Lewthwaite, personal communication). The participants were asked about their level of confidence or certainty in performing a given task using their contralesional hand alone or in conjunction with the ipsilesional hand (e.g. “How confident are you that with your [L/R] hand you can cut food with a knife and fork at a restaurant?”). Each item is scored from 0 to 100, with 100 indicating a very high level of confidence or certainty. Scores are averaged across the 20 items to render a Total score between 0 and 100. In a previous study,^9^ the CAHM was found to be both valid and reliable (Cronbach’ alpha = 0.96 and ICC = 0.91).

### Amount of Use Test (AAUT)

The Actual Amount of Use Test (AAUT) was developed by Taub, Crago, and DeLuca as an assessment of non-use.^15,30,31^ In this context, non-use is defined as the difference between the ability to perform a task with the contralesional hand and the choice to use that hand. Participants perform 14 upper extremity tasks, first in a “spontaneous” condition, i.e., without any instruction, supervision or time limits, and next in a “forced” condition ^32^, wherein they are expressly instructed to use their contralesional hand. Video data are recorded in each condition. For the spontaneous condition, video recording is conducted unbeknownst to the participant, who is debriefed at the end of the test.

Video data were analyzed post-hoc by a trained observer in the Winstein Lab to assess two primary measures: amount of use (AOU) and Quality of Movement (QOM). The AOU is defined as the choice to use the contralesional hand and is coded as 0 or 1, where 0 indicates that the contralesional hand was not used. Thus, a maximum total of 14 points is possible. The QOM is an index of the quality of movement performance and is quantified for each item using a 6-point Likert scale (0 to 5), where 0 is a non-attempt and 5 indicates successful and fairly quick completion of the task using the contralesional hand. A QOM score of 1 indicates an unsuccessful attempt, whereas a score of 2, 3 and 4 indicate purposeful attempts that are either influenced by abnormal movement synergies or are slow or inaccurate. The sum of QOM scores therefore ranges between 0 and 70.

The sums of AOU and QOM were quantified for the spontaneous conditions (sAOU and sQOM) as well as for the forced conditions (fAOU and fQOM). Non-use (NU) was defined as the relative difference between AOU for the spontaneous and forced conditions (normalized for performance in the forced condition):

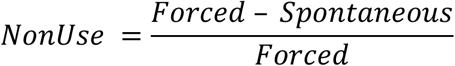

### Data analysis approach

Prior to statistical analyses, all data were inspected and distributions not meeting normality assumptions were appropriately transformed; positively skewed distributions (Education, sAOU) were square root transformed and negatively skewed distributions (fAOU, sQOM, and fQOM) were square transformed.

We assessed whether AAUT indices differed as a function of hemisphere of stroke using two sample t-tests. The relationship of AAUT indices to one another was assessed with Pearson correlation tests. We used two complimentary data analysis approaches to understanding the relationship of predictor variables to AAUT scores. We first determined the individual relationships of demographic predictors and test battery scores to AAUT scores with Pearson correlations. We used a lenient threshold for significance (i.e., p < .05 uncorrected for multiple comparisons) in the service of the next stage of data analysis, in which we determined which of the significant individual indices contributed to predicting AAUT scores when the co-variance among the predictors was taken into consideration. Accordingly, we built multiple regression models with the AAUT indices as dependent measures and UEFM scores and any measure that demonstrated significant individual correlations with AAUT scores as predictor variables. For each set of models, we then implemented a backward stepwise regression procedure wherein we used an r^2^ change test to compare the improvement in fit of the full model versus a model in which the least predictive variable was removed. Finally, given that the UEFM is correlated with most of the background variables, entering it in the model reduces our ability to understand the degree to which the background tasks jointly predict NU when just the relationships between those tasks is taken into account. Therefore, a final multiple regression model examined which of the background test indices contributed to predicting NU when UEFM was *not* entered.

## Results

Table 2 shows scores on the background battery and AAUT.

**Table 2.** AAUT and background test performance.

| Subject | sAOU | fAOU | sQOM | fQOM | NU | VRLAT:<br>Attention/<br>Arousal | VRLAT:<br>Neglect | CAHM | Limb<br>Apraxia<br>(% cor.) |
| --- | --- | --- | --- | --- | --- | --- | --- | --- | --- |
| S1 | 2 | 10 | 0.38 | 0.58 | 0.80 | 0.98 | -0.03 | 36.25 | 0.95 |
| S2 | 3 | 11 | 0.36 | 0.47 | 0.73 | 0.95 | 0.00 | 21.25 | 0.90 |
| S3 | 6 | 14 | 0.90 | 0.83 | 0.57 | 0.90 | 0.00 | 53.00 | 0.72 |
| S4 | 3 | 13 | 0.69 | 0.74 | 0.77 | 0.93 | 0.03 | 43.00 | 0.82 |
| S5 | 2 | 13 | 0.67 | 0.72 | 0.85 | 0.95 | -0.05 | 74.00 | 0.90 |
| S6 | 4 | 14 | 0.98 | 0.97 | 0.71 | 0.90 | 0.00 | 61.75 | 0.77 |
| S7 | 7 | 14 | 0.94 | 0.97 | 0.50 | 0.98 | 0.03 | 59.50 | 0.80 |
| S8 | 6 | 14 | 0.98 | 0.97 | 0.57 | 0.98 | 0.03 | 73.00 | 0.90 |
| S9 | 10 | 14 | 1.00 | 0.97 | 0.29 | 1.00 | 0.00 | 90.50 | 0.87 |
| S10 | 6 | 14 | 0.95 | 0.94 | 0.57 | 1.00 | 0.00 | 48.00 | 0.77 |
| S11 | 4 | 14 | 0.46 | 0.46 | 0.71 | 0.98 | -0.03 | 43.05 | 0.77 |
| S12 | 9 | 14 | 0.92 | 0.97 | 0.36 | 0.93 | 0.08 | 54.50 | 0.75 |
| S13 | 3 | 12 | 0.73 | 0.90 | 0.75 | 0.75 | -0.13 | 36.00 | 0.80 |
| S14 | 2 | 13 | 0.55 | 0.60 | 0.85 | 0.88 | 0.03 | 52.50 | 0.87 |
| S15 | 6 | 14 | 0.70 | 0.81 | 0.57 | 0.88 | -0.09 | 81.50 | 0.85 |
| S16 | 2 | 9 | 0.47 | 0.73 | 0.78 | 0.85 | 0.00 | 36.50 | 0.82 |
| S17 | 3 | 12 | 0.64 | 0.77 | 0.75 | 0.88 | 0.03 | 54.50 | 0.82 |
| S18 | 11 | 14 | 0.27 | 0.99 | 0.21 | 1.00 | 0.00 | 74.50 | 0.90 |
| S19 | 3 | 11 | 0.71 | 0.96 | 0.73 | 0.93 | 0.03 | 40.00 | 0.82 |
| S20 | 6 | 14 | 0.85 | 0.89 | 0.57 | 1.00 | 0.00 | 64.25 | 0.90 |

Abbreviations: sAOU, spontaneous amount of use; fAOU, forced amount of use; sQOM, spontaneous quality of movement; fQOM, forced quality of movement; NU, non-use; VRLAT, virtual reality lateralized attention test; CAHM, confidence in arm and hand movement.

### Individual predictors of AAUT scores

#### Demographic and sensory-motor predictors

Participants with right and left hemisphere stroke did not differ in any of the AAUT indices (all t values < 1.36, p values > .18). Further, none of the AAUT indices (sAOU, fAOU, NU, sQOM, fQOM) correlated with age, chronicity, or education (all r values ≤ .40, all p values > .05).

Table 3 shows that all AAUT indices were significantly correlated with each other (all p values < .05). All of the AAUT indices also correlated with the UEFM (p’s < .01).

**Table 3.**
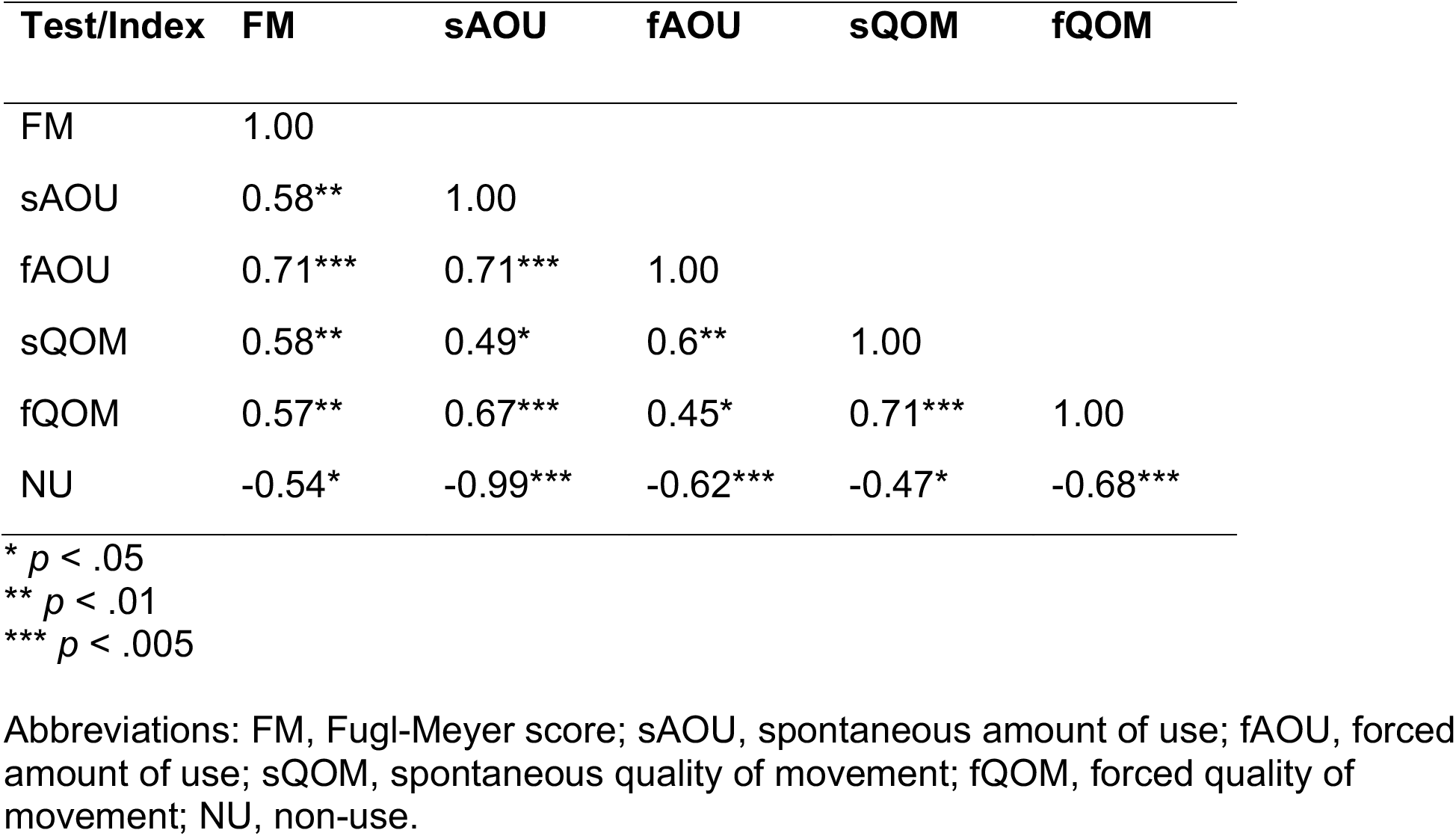
Pearson correlation coefficients for AAUT indices

Although overall stroke severity is likely to substantially drive these correlations, Figure 1A shows that 40% (8/20) of participants have mild functional impairment (UEFM > 45) but nevertheless spontaneously use their contralesional arm < 50% of the time. Moreover, as Figure 1B shows, numerous individuals who spontaneously use their arm < 50% of the time have QOM scores > 80%, indicating purposeful albeit slightly slower or less accurate movements. The data thus far suggest, therefore, that factors other than overall severity may influence the results.

**Figure 1.**
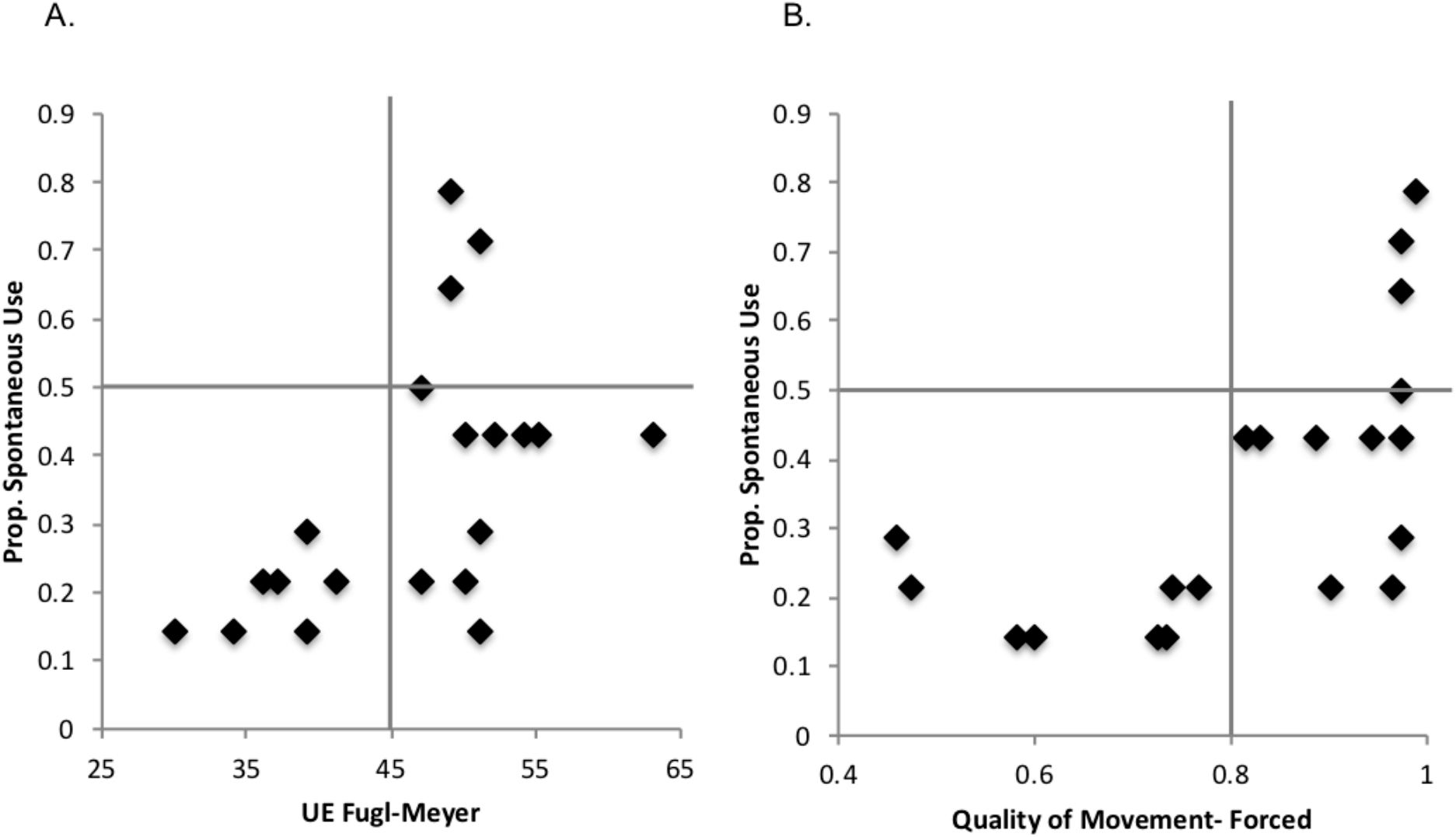
A) Relationship between proportion of spontaneous use in the AAUT and upper extremity Fugl-Meyer. **B)** Relationsh ip between proportion of spontaneous use in the AAUT and quality of movement in the AAUT.

#### Psychological and neuropsychological predictors of non-use

CAHM was significantly correlated with all of the AAUT indices (sAOU: r = .61, p =.004; fAOU: r = .67, p = .001; NU: r =-.55, p = .01; sQOM: r =.45, p = .045; fQOM: r =.52, p = .02. Thus, greater confidence in movement was associated with a higher amount of use and quality of movement, and critically, reduced non-use.

Lower total scores on the VRLAT (attention/arousal) were associated with decreased sAOU (r = .47, p = .04) and increased NU (r = -.45, p = .04). The relationship between VRLAT total scores and fAOU was not significant (r =.38, p = 0.1).

The normalized difference score from the VRLAT (spatial neglect) did not predict any of the AAUT indices (all r’s < .20, p’s > .40).

There was a trend for correlation of the limb apraxia score with sQOM (r = -.41, p = .07), but in a direction opposite to that predicted (higher quality of movement, worse apraxia score).

### Combined predictive models of AAUT indices

Using backward stepwise regressions with AAUT scores as dependent measures, we sought the simplest predictive models that resulted in an insignificant deterioration in model fit as compared to the next most complex models. Results are shown in Table 4. Not surprisingly, the full models (i.e., including UEFM and other indices that were individually correlated with the dependent measure) significantly predicted each of the dependent measures.

**Table 4.**
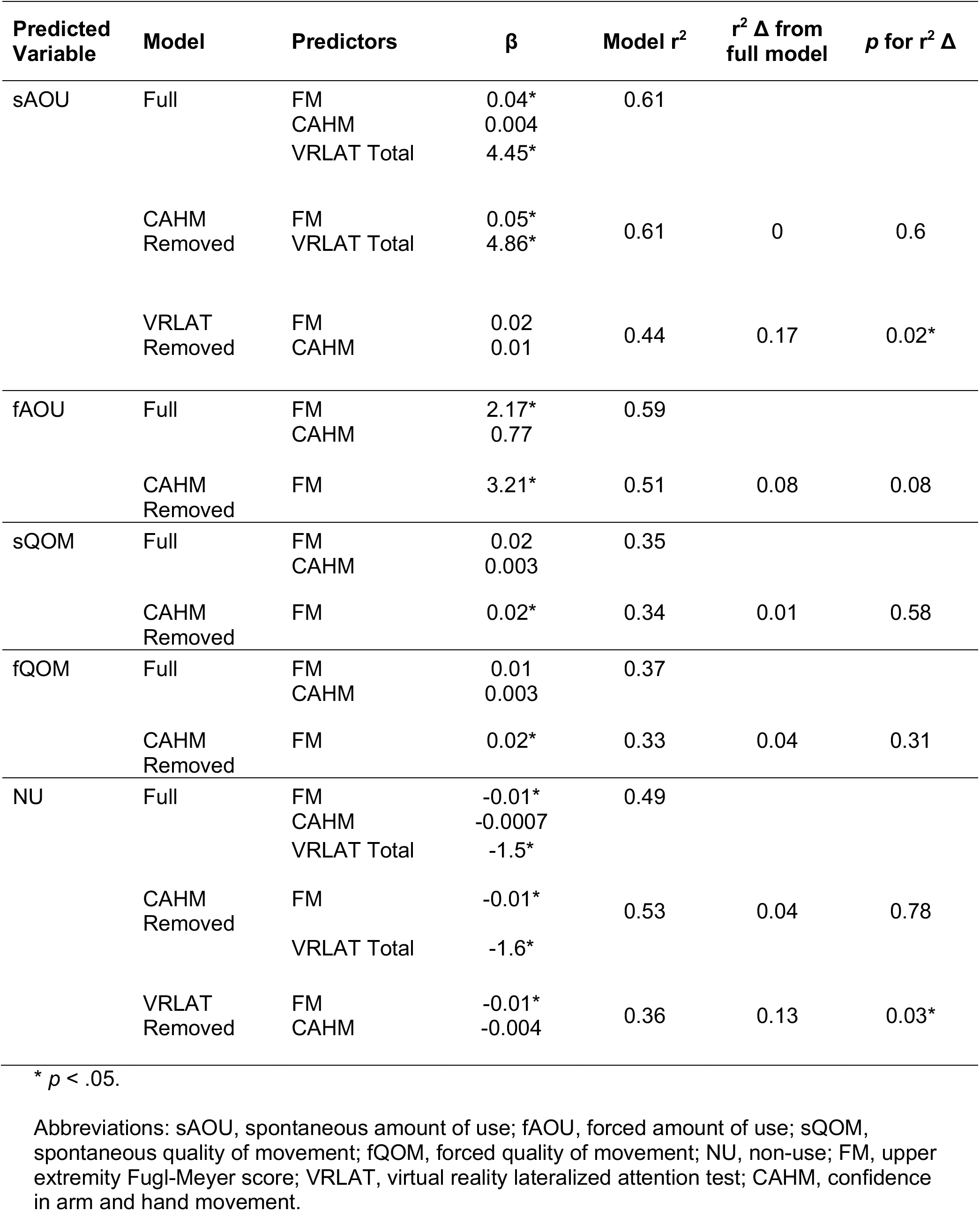
Results of step-wise regression models.

| Predicted Variable | Model | Predictors | $\beta$ | Model $r^2$ | $r^2 \Delta$ from full model | $p$ for $r^2 \Delta$ |
| --- | --- | --- | --- | --- | --- | --- |
| sAOU | Full | FM<br>CAHM<br>VRLAT Total | 0.04*<br>0.004<br>4.45* | 0.61 |  |  |
|  | CAHM Removed | FM<br>VRLAT Total | 0.05*<br>4.86* | 0.61 | 0 | 0.6 |
|  | VRLAT Removed | FM<br>CAHM | 0.02<br>0.01 | 0.44 | 0.17 | 0.02* |
| fAOU | Full | FM<br>CAHM | 2.17*<br>0.77 | 0.59 |  |  |
|  | CAHM Removed | FM | 3.21* | 0.51 | 0.08 | 0.08 |
| sQOM | Full | FM<br>CAHM | 0.02<br>0.003 | 0.35 |  |  |
|  | CAHM Removed | FM | 0.02* | 0.34 | 0.01 | 0.58 |
| fQOM | Full | FM<br>CAHM | 0.01<br>0.003 | 0.37 |  |  |
|  | CAHM Removed | FM | 0.02* | 0.33 | 0.04 | 0.31 |
| NU | Full | FM<br>CAHM<br>VRLAT Total | -0.01*<br>-0.0007<br>-1.5* | 0.49 |  |  |
|  | CAHM Removed | FM<br>VRLAT Total | -0.01*<br>-1.6* | 0.53 | 0.04 | 0.78 |
|  | VRLAT Removed | FM<br>CAHM | -0.01*<br>-0.004 | 0.36 | 0.13 | 0.03* |

\* $p < .05$ .
Abbreviations: sAOU, spontaneous amount of use; fAOU, forced amount of use; sQOM, spontaneous quality of movement; fQOM, forced quality of movement; NU, non-use; FM, upper extremity Fugl-Meyer score; VRLAT, virtual reality lateralized attention test; CAHM, confidence in arm and hand movement.

For both sAOU and NU, reduced models with just UEFM and VRLAT total included (i.e., with CAHM removed) accounted for as much variance in predicting the dependent measures as the full models with CAHM included. However, removal of the VRLAT resulted in a more poorly-fitting model as compared to the models with the VRLAT included. Thus, the preferred model for predicting sAOU and NU contains UEFM and VRLAT total. For fAOU, the best fitting model included both UEFM and, at a trend level, CAHM; conversely, removal of CAHM resulted in a model that tended to fit more poorly.

For both sQOM and fQOM, the simplest predictive models contained UEFM alone.

To address neuropsychological and psychological predictors of NU irrespective of sensory-motor impairment, we ran a final model in which UEFM was not considered. Based on their significant individual correlations with the AAUT indices, CAHM and VRLAT Total scores were entered as independent variables (adjusted r^2^ = .32, F (2,17) = 5.4, p < .05). Only CAHM emerged as a significant predictor of NU in this model (CAHM β = -.005, SE = .002, t = −2.27, p = .04; VRLAT β = -.98, SE = .63, p = .14).

## Discussion

This study provides preliminary evidence relevant to the demographic, sensory-motor, psychological and neuropsychological factors predicting arm non-use in a sample of 20 chronic stroke survivors, half of whom had left- and half right-hemisphere lesions. We observed, first, that approximately 40% of patients with mild impairment on the upper extremity Fugl-Meyer (UEFM) nevertheless used their contralesional limb spontaneously less than half the time, broadly consistent with prior reports of NU frequency.^33^ We showed, further, that non-use, as indexed by the disparity between spontaneous and forced limb use, was not related to demographic factors or hemisphere of lesion in this sample. On the other hand, scores on a common measure of sensory-motor impairment, the UEFM, predicted the degree of non-use, and also predicted spontaneous amount of use and quality of movement. Thus, patients with greater sensory-motor impairment showed a larger relative disparity between their arm use in the forced use condition and the spontaneous use condition, again consistent with prior observations.^7,33^ In other words, the disparity between capability and use is not monotonic, suggesting that non-use is a particularly important issue in moderately-impaired (as compared to mildly-impaired) individuals. Importantly, extending prior observations, we demonstrated that non-lateralized attention/arousal predicted both spontaneous use and non-use (but not forced use) above and beyond sensory-motor impairment. When considering all individual psychological and neuropsychological predictors of NU (without also considering their correlations with UEFM), an index of self-efficacy (confidence in one’s own hand and arm movement) predicted non-use. Thus, depending on the measures that are considered together, both psychological and neuropsychological factors significantly contribute to the prediction of non-use. We will discuss these results in turn.

### Attention/arousal as a predictor of non-use

The observed relationship between non-lateralized attention and non-use is consistent with prior observations regarding the relationship of attention and motivation. Non-lateralized attention/arousal was classically described as relying on ascending pathways originating from the mid-brain reticular formation.^34^ More recent research has identified multiple circuits critical to attention and arousal, involving thalamic nuclei, hypothalamus, limbic regions, cingulate, frontal operculum, and basal ganglia. Accordingly, even small subcortical strokes may be associated with deficits in vigilance and attention.^35^ Ascending pathways through subcortical structures are modulated by a number of descending inputs from cortical regions including frontal and parietal cortex. ^36^ Deficits in non-lateralized attention, though potentially more common after right hemisphere stroke ^e.g.,37^ are observed in strokes affecting either hemisphere.^e.g.,38^ Recently, Rinne et al. ^39^ demonstrated in both neurotypical aged and hemiparetic stroke participants a relationship between motor performance (dexterity and strength) and attention control, as measured by distractor resistance, even controlling for lesion size and baseline performance.

Previous investigations suggest that non-lateralized attention/arousal bears a relationship to mental effort and to goal-driven aspects of behavior. For example, arousal/effort as measured by pupil dilation appears necessary to direct eye gaze toward objects that are not salient but nevertheless relevant to current goals. Conversely, low arousal/effort as assessed by pupil constriction is associated with attentional capture by highly salient but irrelevant stimuli.^40^ An even more direct link between attention and goal-driven behavior is suggested by studies showing that individuals high on scales of “grit” – the ability to persevere toward long term goals— demonstrate greater and more efficient sustained attention in laboratory tasks.^41,42^

It is likely that using even a mildly paretic arm requires effort and motivation.^16^ Addressing this possibility, several clinical trials are assessing whether improving motivation may impact stroke rehabilitation outcome. For example, the ongoing WAVES trial ^43^ and others^44^ are assessing the effects of accelerometers that track arm use and provide feedback to improve motivation, and the Armeo Senso Reward trial will compare game feedback and contingent monetary rewards versus non-rewarded arm training.^45^ The successful Queen’s Square Program ^46^ includes motivating computer games, self-efficacy, and goal-setting in addition to intensive active practice. Focus groups conducted after the program indicated that patients believed that individualized goals and motivation were key ingredients in the program’s success. Finally, our group recently reported results of a planned intention-to-treat analysis of a large-scale study comparing an Accelerated Skill Acquisition Program (ASAP) that included motivational enhancements, autonomy support, and critical elements of the transfer package (e.g. promotion of self-efficacy) with dose-matched usual therapy. Importantly, we demonstrated that the ASAP intervention substantially accelerated improvements across a spectrum of patient reported outcomes that included physical function, reintegration into normal living and health-related quality of life, and exacted lasting gains in patient-reported overall strength.^47^

### Self-efficacy as a predictor of non-use

The individual correlations we performed suggested that greater confidence in arm and hand movement was associated with greater spontaneous amount of use and quality of movement, and reduced non-use. These relationships were weakened when we considered sensory-motor impairment in the same model, reflecting the shared variance between perceived confidence in movement and sensory-motor impairment. In other words, patients’ perceived confidence in what they could achieve in the future was realistically influenced by their abilities, and the latter was a relatively robust predictor of non-use. On the other hand, when we considered psychological and neuropsychological predictors of non-use alone (i.e., non-lateralized attention and confidence in arm and hand movement), only the latter demonstrated a robust relationship with non-use. Though caution is required in interpreting these results, the data suggest that self-efficacy may (weakly) contribute to the prediction of non-use and should be further evaluated in future studies.

Recent evidence suggests that reduced self-efficacy may be associated with poorer rehabilitation outcomes. For example, a recent study with 120 patients showed that initial values of the Generalized Self-Efficacy Scale were correlated with scores on the Barthel Index and Rivermead Mobility Index (and other measures) performed after 3 weeks of daily rehabilitation treatment. ^48^ Following from such data, recent approaches to stroke rehabilitation have increasingly emphasized the importance of interventions targeting self-efficacy. Sit et al. ^49^, for example, performed a two arm single-blind randomized controlled trial of 210 stroke patients and demonstrated that an “empowerment” intervention that included goal-setting and action planning improved outpatient rehabilitation outcomes as compared to treatment without the added intervention (see also ^50^). Indeed, a recent phase IIb RCT of therapy dose in chronic stroke survivors that used the ASAP intervention demonstrated a significant effect of dose on a patient-reported outcome measure of arm use (i.e. Motor Activity Log-Quality of Movement), supporting the importance of perceived self-efficacy for effecting gains in spontaneous arm use in the natural environment ^51,52^.

### Study limitations

Contrary to our expectation, we failed to observe a relationship between either limb apraxia or spatial neglect and limb non-use. Nor did we observe a relationship between variables such as age or hemisphere of stroke that might plausibly contribute. Due to the small sample size, caution should be exercised in interpreting these and any other null results. In addition, our selection criteria, which required Fugl-Meyer scores in the range of mild to moderate impairment, may have reduced our ability to observe some of these relationships. Specifically, because both apraxia and neglect severity tend to co-vary with overall stroke severity, our selection criteria appear to have resulted in exclusion of patients with moderate to severe neglect or limb apraxia. Given that (as noted earlier) both disorders are associated with poor rehabilitation outcomes, it remains possible that they may contribute to the prediction of limb non-use in more severely impaired patients. Additional studies with a larger sample and broader range of patient severity will be necessary to clarify these relationships.

The sample in this preliminary study was limited in several other potentially important respects. No information was available about medical co-morbidities that may influence spontaneous arm use, such as peripheral neuropathy. We also did not have information about whether stroke was of ischemic, hemorrhagic or embolic origin; stroke type has been shown to influence outcomes overall, and it is unclear whether it may affect non-use.^53^ Additionally, we did not assess overall cognitive function, pain, or mental health. Pain of the contralesional limbs has been hypothesized to be associated with non-use.^1^ In terms of mental health, depression, in particular, has been negatively associated with rehabilitation outcome.^e.g.,54^ Of interest for future research is the question of whether depression may mediate the relationships we observed between attention/arousal and non-use, and confidence in movement and non-use.

## Conclusion

In this preliminary study we demonstrated that psychological and neuropsychological factors contribute to the prediction of upper extremity non-use in chronic stroke survivors with mild-to-moderate sensory-motor impairments. Given the complexity of the non-use phenomenon, it follows that a combination of impairment, attention/arousal, and perceived confidence predicted non-use behavior. That a measure of attention/arousal predicted non-use behavior above and beyond sensory-motor functioning highlights the importance of motivated engagement to drive use of the paretic limb. As such, it is increasingly clear that rehabilitation efforts are more likely to be successful when they engage the participant by providing meaningful task practice and motivational enhancements (see ^55^ for discussion). Larger-scale studies of the non-use phenomenon in individuals ranging in severity, and with incorporation of additional measures (e.g., mental health, lesion volume and white matter connectivity, pain, trait- and state-level motivation) will be required to extend these findings.

## Acknowledgments

This research was supported by the Moss Rehabilitation Research Institute and the Division of Biokinesiology and Physical Therapy. We appreciate the participation of the individuals whose data are included in this study.

The authors declare no conflicts of interest.

